# Identifying the *in vivo* cellular correlates of antipsychotic drugs

**DOI:** 10.1101/226050

**Authors:** Radhika S. Joshi, Mitradas M. Panicker

## Abstract

Antipsychotics have revolutionized the treatment of mental illness from the 1950s (Hippius, 1989; Shen, 1992). Even today antipsychotics are the preferred treatment for a number of mood disorders including schizophrenia, bipolar disorder, obsessive-compulsive disorder, severe depression etc. (Blier, 2005; Cookson, 2008; Leucht et al., 2009; McDougle et al., 2000). However, the mechanism of action of antipsychotics still remains sketchy and controversial. The brain areas, neural circuits and cellular targets involved in the effects of antipsychotics need to be better identified.

Drug binding studies suggest dopamine receptor D2 and serotonin receptor 5-HT_2A_ as the prime targets of antipsychotics based on binding affinities (Roth et al., 1994, 2004; Yadav et al., 2011a). Based on the relative affinity for the 5-HT_2A_ and D2, antipsychotics are also classified into typical and atypical classes. Typical antipsychotics exhibit higher affinity for D2 than 5-HT_2A_ and the reverse is seen for atypical antipsychotics (Meltzer et al., 1989). Antagonism at the 5-HT_2A_ and D2 is thought to underlie some of the therapeutic effects and/or side effects of antipsychotics in patients and animal models (Fribourg et al., 2011; Kapur et al., 1995, 2000; Moreno et al., 2016; Wadenberg et al., 2000). Along with the 5-HT_2A_ and D2, antipsychotics can also bind to many other GPCRs with varying affinities, for example, muscarinic receptors, adrenergic receptors and histamine receptors (Roth et al., 2004). However, role of these GPCRs in modulating the antipsychotic-induced effects on neural circuits or cellular targets is largely unknown.

Previously *c-fos* activity has been used to identify the brain areas and cells that are active on administration of antipsychotics. Up-regulation of *c-fos* gene is a bonafide marker for neuronal activity in cell culture, rodent and the human brain. Many stimuli have been shown to cause induction of *c-fos*, for example seizure (Gunn et al., 1990; Morgan et al., 1987), membrane depolarization (Sheng et al., 1990), novel environment and psychoactive drugs (Day et al., 2001; Ostrander et al., 2003; Uslaner et al., 2001), etc.

Prior studies showed that typical and atypical antipsychotics cause increased *c-fos* activity in the striatum and prefrontal cortex respectively, whereas up-regulation of *c-fos* in the nucleus accumbens is reported to be activated by both (Deutch A. Y., 1996; Verma et al., 2007; Wan et al., 1995). Although these studies have provided valuable initial information, they have certain shortcomings. In these studies, *c-fos* activity was detected by immunohistochemistry or in-situ hybridization. Thus the sensitivity of the antibody and the probe as well as limited access to the morphology of the cells limits the characterizations. Importantly the information is visualized in fixed tissue, severely restricting the availability of the stimulus/antipsychotic responsive cells for further biochemical or physiological investigation.

In this study we describe the activity induced by antipsychotics, using the ‘FosTrap’ system. This system designed by Luo and colleagues (Guenthner et al., 2013) permanently marks cells in which *c-fos* is active if also provided with Tamoxifen. We used F1 progeny of FosCreER^T2^ mice and Lox-tdTomato mice for our experiments. Here *c-fos* promoter drives the expression of CreER^T2^ recombinase. The presence of ER site allows entry of Cre recombinase into the nucleus only in the presence of Tamoxifen. Therefore Tamoxifen, coupled with the stimulus, traps the ‘stimulus responsive cells’ to permanently express tdTomato (Guenthner et al., 2013) (Figure 1a). Since the tdTomato is cytoplasmic, the entire cell can be visualized even without any fixation or processing. ‘FosTrap’ mice have shown increased *c-fos* activity following exposure to novel context, whisker stimulation, light exposure, etc. which led us to try and identify cells that are activated by antipsychotics in the mouse brain (Guenthner et al., 2013).

**Figure 1.**
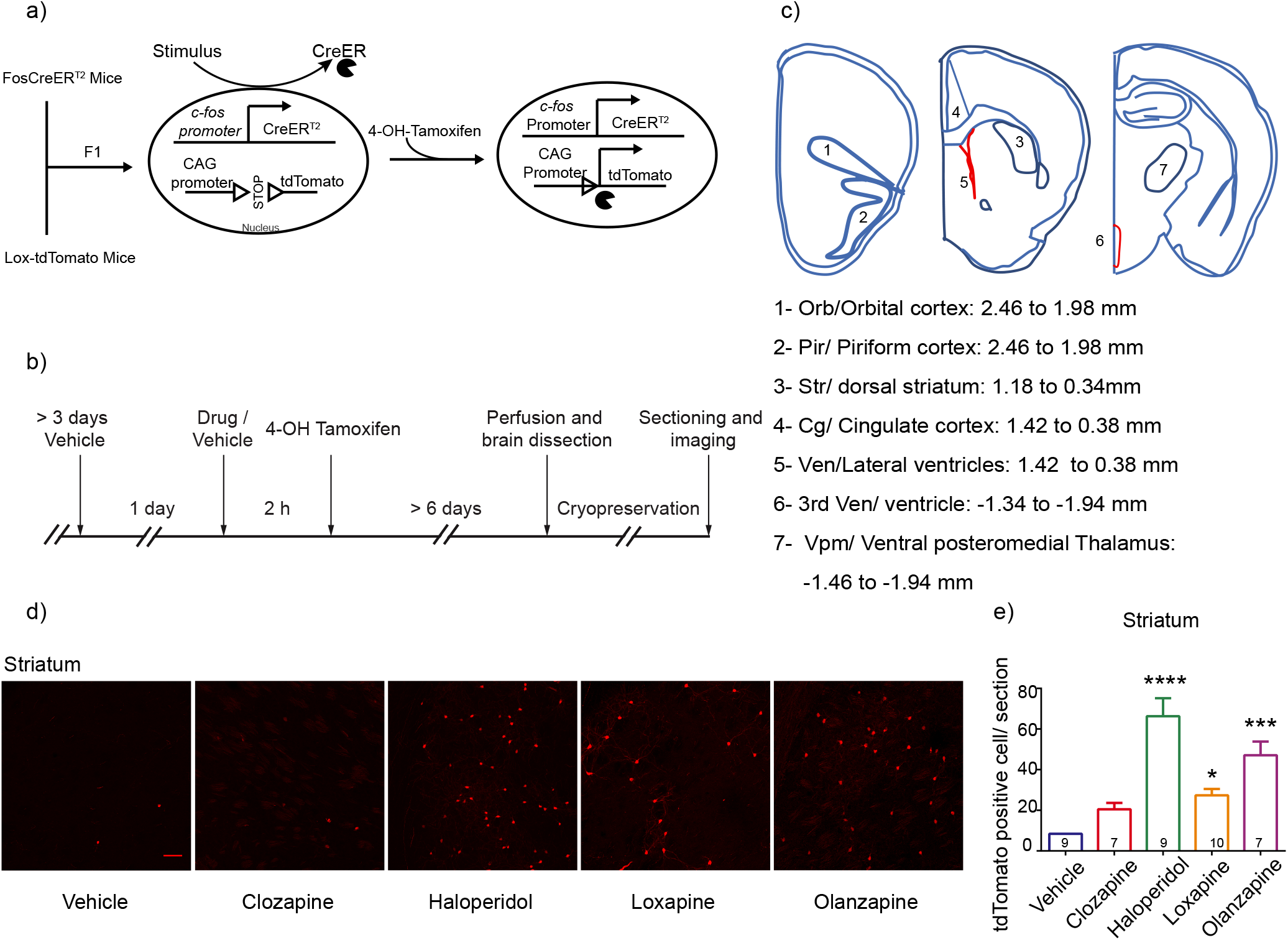
FosTRAP mice show increased c-Fos labelling in striatum on treatment with antipsychotic drugs. a) Schematic representation of the ‘FosTRAP’ system. F1 progeny of FosCreER and Lox-tdTomato mice was used for experiments with the genotype *Fos^CreER/+^ R26^Al14/+^*. In the presence of a stimulus, responsive neurons drive expression of Cre recombinase. Cre recombinase protein cannot enter the nucleus until bound by Tamoxifen or 4-OH-Tamoxifen. Once bound by 4-OH-Tamoxifen Cre enters the nucleus and drives the expression of a fluorescent protein, in our case tdTomato. Then on, the cell is genetically committed to express tdTomato. Arrowheads inside the nucleus indicate initiation of transcription. Solid sector denotes CreER^T2^. b) Schematic of the experimental protocol. Antipsychotic drugs or vehicle were used as a stimulus. 4-OH-Tamoxifen was administered 2 hours later. The mice were perfused minimum of 6 days later and the brain slices were sectioned. c) Schematic representation of the areas imaged. The region of interest was observed with the help of Hoechst stain. d) Representative images of the tdTomato labelling in dorsolateral striatum. Scale Bar-10 μM. e) Quantification of the number of tdTomato positive cells per section. Haloperidol showed the maximum number of tdTomato positive cells in the striatum, followed by Olanzapine and Loxapine. Clozapine showed a trend towards increased number of tdTomato positive cells compared to vehicle. Numbers in the bars represent the number of mice in that group. Drug and vehicle groups were compared by Kruskal-Wallis test with Dunn’s multiple comparison. Data represented as Mean ± SEM. * p< 0.05, ** p<0.01, *** p< 0.001, **** p< 0.0001.

We tested four different antipsychotics from typical and atypical groups viz., Clozapine, Haloperidol, Loxapine and Olanzapine. 5-HT_2A_ being the primary target of Clozapine, we also investigated the role of 5-HT_2A_ in modulating the activity pattern induced by Clozapine. Using this system we screened different cortical and subcortical brain areas for active *c-fos* in response to antipsychotics. In general, we observed that the typical and atypical antipsychotics showed little overlap in their pattern of activity. The brain areas such orbital cortex, piriform cortex and ventral-posteromedial thalamus (Vpm) were responsive to the atypical drugs. Importantly, we identified ependymal cells within the ventricles as a novel cellular target of the antipsychotic Clozapine and Olanzapine, and these were heavily modulated by 5-HT_2A_. Briefly, the ‘FosTrap’ system has allowed us to identify brain regions and cell types which are known to be activated by antipsychotics as well as newer regions and cell types. Since the TRAP system allows easy and direct visualization of the antipsychotic responsive cells it can be used further for potential manipulation and investigation of the trapped cells *in vivo*.

## Material and Methods

### Animals

Animals were maintained on ad libitum food and water on a day-night cycle of 10-14 hours. Experiments were performed during the daytime. Males and females, minimum of 8 weeks or older were used for the experiments. FosCreER mice (c-Fos Cre ERT2 (B6.129(Cg)-*Fos^tm1.1(cre/ERT2)Luo^*/J) and Lox-tdTomato (B6.Cg-*Gt*(*ROSA*)*26Sor^tm14(CAG-tdTomato)Hze^*/J) mice were obtained from the Jackson laboratory, (Stock No: 021882 and stock No: 007914, respectively) and maintained as per the Jackson laboratory guidelines. *Htr2a^−/−^* mice were generated in-house and maintained by heterozygous matings as described previously (Joshi et al., 2016). Animal usage protocols were approved by the Institutional Animal Ethics Committee.

F1 progeny of cross between FosCreER^T2^ mice (*Fos^CreER/+^*) and Lox-tdTomato [*R26^Al14/A14^*) was used for experiments. The FI (*Fos^CreER/+^ R26^Al14/+^*) mice were crossed further into *Htr2a^−/−^* background to generate the triple transgenic mice with genotype *Fos^CreER/+^ R26^Al14/+^Htr2a^+/+^* and *Fos^CreER/+^ R26^Al14/+^Htr2a^−/−^*. The *Fos^CreER/+^ R26^Al14/+^ Htr2a^−/−^* were maintained on the *Htr2a^−/−^* background. Similarly the *Fos^CreER/+^ R26^A114/+^ Htr2a^+/+^* mice were maintained on the *Htr2a^+/+^* background. To rule out the effect of maternal rearing, experiments were conducted on a small set of *Htr2a^+/+^* and *Htr2a^−/−^* animals obtained by heterozygous matings of *Fos^CreER/+^ R26^Al14/+^ Htr2a^+/−^*.

All the mice were genotyped for the presence of Cre and Lox locus before use to achieve *Fos^CreER/+^ R26^Al14/+^*. If required mice were genotyped for the *Htr2a* locus as described previously (Joshi et al., 2016).

#### Genotyping

Genotyping was performed by standard polymerase chain reaction. The genomic DNA was obtained from tail biopsies.

##### Primers

For Htr2a knockout strain, Wild Type (WT) product: 408bp, Htr2a Mutant product: 642 bp WT Forward-CAT GGA AAT TCT CTG TGA AGA CA, WT Reverse-AGG ATG GTT AAC ATG GAC ACG, Mutant Forward-AGT TAT TAG GTC CCT CGA AGA GGT, Mutant Reverse-GGT ACA AGT CCT TGC TGT ACA ATG.

FosCreER mice, WT product: 215, Mutant product: 293

Common Forward-CAC CAG TGT CTA CCC CTG GA, WT Reverse-CGG CTA CAC AAA GCC AAA CT, Mutant Reverse-CGC GCC TGA AGA TAT AGA AGA.

For Lox-tdTomato mice, WT product: 297 bp, Mutant product: 196 bp

WT Forward-AAG GGA GCT GCA GTG GAG TA, WT Reverse-CCG AAA ATC TGT GGG AAG TC, Mutant Reverse-GGC ATT AAA GCA GCG TAT CC, Mutant Forward-CTG TTC CTG TAC GGC ATG G.

#### Drugs

Clozapine (Catalogue No. 0444) stock: 50mg/ml, Haloperidol (Catalogue No. 0931) stock: 10mg/ml, Olanzapine (Catalogue No. 4349) stock: 50mg/ml, were from Tocris Bioscience, (Bristol, UK) and dissolved in DMSO. Loxapine (Catalogue No. L106) stock: 10mg/ml was from, Sigma-Aldrich, USA and dissolved in 0.9% saline. All aqueous solutions were buffered to pH 6 – 6.5 if required. The drugs were administered intraperitoneally.

4-OH-Tamoxifen (Catalogue No. H6278) was purchased from Sigma-Aldrich, USA. It was prepared as described previously (Guenthner et al., 2013). Briefly, 4-OH-Tamoxifen was dissolved in ethanol at 20mg/ml and stored at −20°C. 4-OH-Tamoxifen stock in ethanol was mixed with corn oil to achieve the final concentration of 10 mg/ml. The ethanol was evaporated using Centrivap before injecting the mice.

#### Mouse brain processing

The animals were perfused with 4% paraformaldehyde (PFA) and the brains were dissected out. Brains were post-fixed in 4% PFA overnight. Following cryopreservation in 30% sucrose, the brains were sectioned into 40μm thick slices.

#### Antibody staining

Anti Vimentin antibody (Catalogue No: ab92547 from Abcam (Cambridge, United Kingdom)) and anti S100 beta antibody (Catalogue No: Z0311 from Dako (Agilent Technologies, Santa Clara, California, USA)) were used for staining. 0.3% Triton in 3% milk powder, prepared in PBS, was used as blocking. The sections were incubated with the primary antibody Vimentin (1:300) and S100β (1:500) overnight at 4°C, followed by staining with the secondary antibodies.

#### Image acquisition and analysis

The brain slices were imaged on Olympus FV1000 (Melville, NY, USA), confocal microscope. Every 3^rd^ section was imaged for Orbital cortex (Bregma 2.46 to 1.98 mm), Piriform cortex (Bregma 2.46 to 1.98), Vpm (Bregma −1.46 to −1.94 mm) and the 3^rd^ Ventricle (Bregma −1.34 to −1.94 mm). Every 6^th^ slice was imaged for the cingulate cortex (Bregma 1.42 to 0.38 mm), dorsal striatum (caudate-putamen) (Bregma 1.18 to 0.34 mm) and lateral ventricles (Bregma 1.42 to 0.38 mm). The areas of interest were identified with the help of Hoechst stain and The Mouse Brain atlas, Franklin and Paxinos. Images were processed using ImageJ 1.47, (NIH, Bethesda, Maryland, USA) software and the number of cells in each image were counted using Cell profiler (Openware, Broad Institute Imaging Platform, Cambridge, Massachusetts, USA). Final data was represented as the number of cells per section averaged from all the slices imaged from the (right and left hemisphere) the area of interest. The ependymal cells were often very close together to count the individual number of cells. Therefore total intensity was counted per image and data was represented as Unit intensity/section.

#### Statistics

Data is represented as Mean ± SEM. Comparison between vehicle and drug treatments were interpreted using one-way ANOVA (or Kruskal Wallis test where appropriate) with correction for multiple comparisons. *Htr2a^+/+^* and *Htr2a^−/−^* genotypes and treatments were compared using two-way ANOVA with correction for multiple comparisons. Student’s t-test was used for comparison between male and female data. The appropriate tests are indicated in the figure legends.

## Results

### Increased *c-fos* activity was observed following treatment with antipsychotics in various brain regions

To identify the antipsychotic responsive cells with a fluorescent marker protein we crossed FosCreER^T2^ mice (B6.129(Cg)-*Fos^tm1.1(cre/ERT2)Luo^*/J) with Lox-tdTomato (B6.Cg-*Gt*(*ROSA*)*26Sor^tm14(CAG-tdTomato)Hze^*/J) and used the FI progeny (*Fos^CreER/+^ R26^Al14/+^* mice) for the experiments (Figure 1b). Following the acute antipsychotic treatment tdTomato positive cells were seen in various brain areas (Figure 1c). The majority of cells that were labelled looked neuronal by morphology. We have used the term-‘tdTomato positive labelling’ interchangeably with *c-fos* activity throughout.

Previous reports show that typical antipsychotics such as Haloperidol cause activation of *c-fos* in the dorsolateral striatum, therefore we first analysed tdTomato labelling in the striatum to validate our system. Dorsolateral striatum showed the highest number of tdTomato positive cells on treatment with Haloperidol. Loxapine, another typical antipsychotic, showed lesser activity in the striatum compared to Haloperidol (figure 1d, e). Surprisingly, Olanzapine which belongs to the atypical class of antipsychotics showed a strong *c-fos* response in the striatum (5.5 fold over control) compared to its structural analogue Clozapine (Figure 1d & e).

In conclusion, the Trap system could successfully and permanently label cells within specific brain regions in an antipsychotic-dependent manner. The areas in which these cells were located were similar to what was expected of the typical antipsychotics based on previous literature. In addition, it has brought out fine differences among the typical and atypical antipsychotics.

### Cortical and thalamic sub-regions are more responsive to Clozapine and Olanzapine

While Clozapine labelled the least number of cells in the striatum, certain cortical regions (Figure 1c) were very responsive to Clozapine. Clozapine and Olanzapine showed *c-fos* activity patterns similar to each other in the cortical structures that we tested, unlike the striatum (Figure 2a, b, c). Briefly, Clozapine and Olanzapine showed significantly higher tdTomato positive cells in the orbital cortex, piriform cortex and anterior cingulate cortex, compared to Vehicle. In these brain areas, the number of tdTomato positive cells induced by Haloperidol and Loxapine was comparable to the Vehicle group.

**Figure 2.**
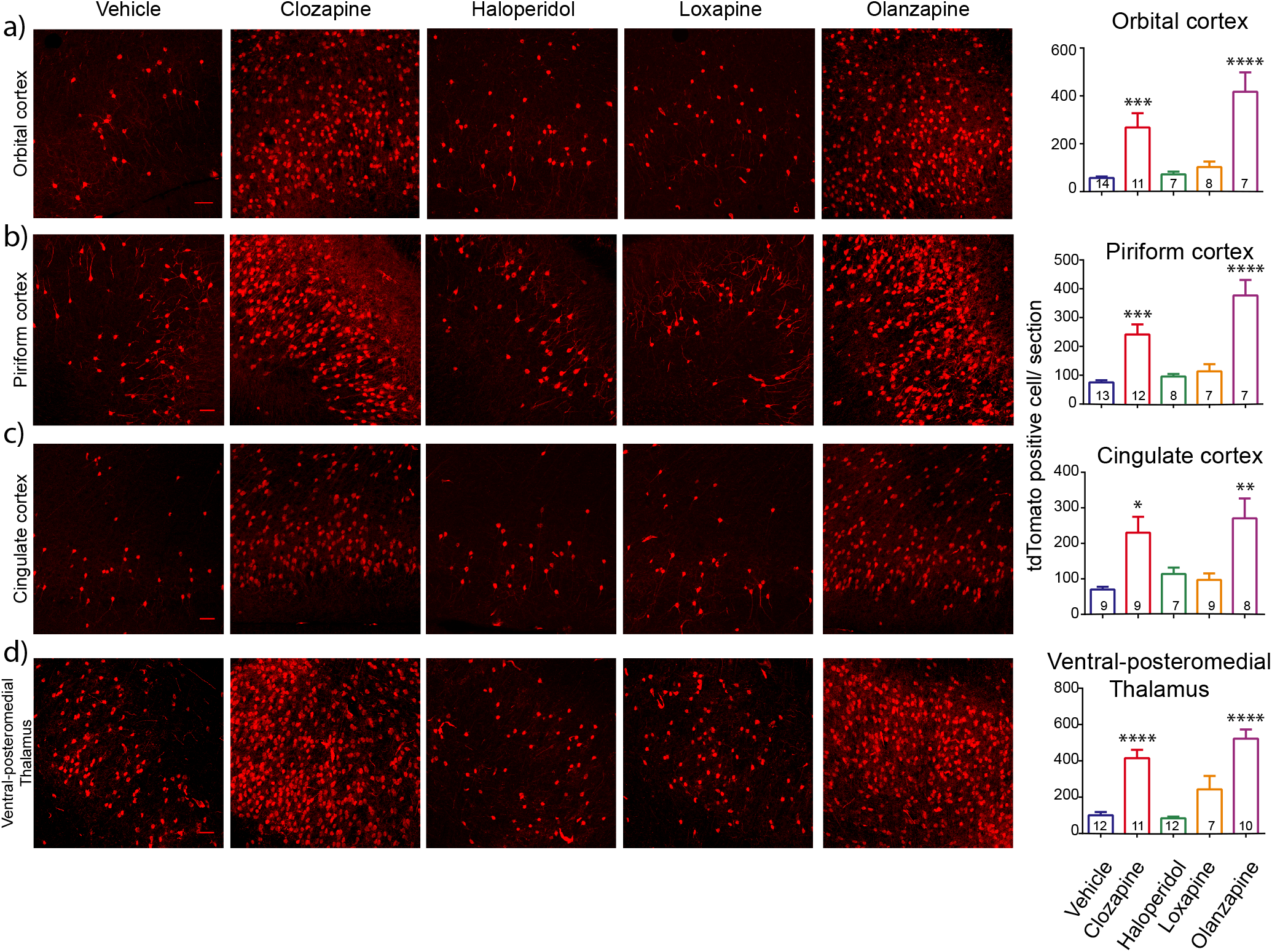
Clozapine and Olanzapine showed increased *c-fos* activity in various cortical regions and Vpm. a- d) Representative images of the tdTomato labelling in orbital cortex, piriform cortex, cingulate cortex and Vpm. Scale Bar-10 μM. The graphs on the right show quantification of the number of tdTomato positive cells per section. In all of these regions Clozapine and Olanzapine showed significantly higher tdTomato positive cells compared to vehicle. The activity induced by Haloperidol and Loxapine was comparable to that of vehicle. Numbers in the bars represent the number of mice in that group. Drug and vehicle groups were compared by one-way ANOVA with Dunne’s multiple comparison and Kruskal-Wallis test with Dunn’s multiple comparison, where appropriate. Data represented as Mean ± SEM. * p< 0.05, ** p<0.01, *** p< 0.001, **** p< 0.0001.

Previous studies have reported higher Clozapine-induced *c-fos* activity in the cortical regions (Deutch A. Y., 1996; Ohashi et al., 2000; Verma et al., 2007), however *c-fos* activity in the orbital cortex and piriform cortex has not been examined in particular.

Along with the cortical regions, *c-fos* activity was noticeable in the Vpm (Figure 1c & 2d). Clozapine and Olanzapine showed higher numbers of tdTomato positive cells in Vpm. Vpm is a part of thalamus which relays oral and facial sensory information to the somatosensory cortex. There are a few reports of antipsychotic-induced *c-fos* activity in the thalamus (Cohen, 1995; Deutch et al., 1995; Rajkumar et al., 2013), although antipsychotic responses have not been reported in Vpm in particular.

Most of the behavioural or biochemical studies have used only male mice, and the responses of female mice remain largely undetermined. Therefore in this study, we also analysed the antipsychotic-induced *c-fos* responses in male as well as female mice. We systematically compared *c-fos* activity in various brain regions between males and females. We did not observe any gender-specific difference in any of the regions tested (Supplementary figure 1a, b) with either Clozapine or Vehicle treatment. Hence females were also included in the subsequent experiments and analysis.

Since antipsychotic effects are dose-dependent, we also analysed a dose response to Clozapine. Previously we had reported significant differences in the Clozapine-induced sedation between *Htr2a^+/+^* and *Htr2a^−/−^* mice at 5mg/kg (Joshi et al., 2016); whereas dose of Clozapine around 20mg/kg has been used very extensively to assess Clozapine-induced *c-fos* responses (Badiani et al., 1999; Deutch A. Y., 1996; Wan et al., 1995). Therefore we tested 0, 5 and 20mg/kg of Clozapine. At 5mg/kg of Clozapine, most brain regions showed a trend towards increased *c-fos* activity (Supplementary figure 1c-f). However at higher dose of 20mg/kg, Clozapine significantly increased the number of tdTomato positive cells in the cortical regions. Doses of the other antipsychotics that we tested were chosen based on the previous literature on behavioural and biochemical effects of these drugs (Cope et al., 2005; Deutch A. Y., 1996; Robertson and Fibiger, 1996).

### Ependymal cells are novel cellular targets of Clozapine and Olanzapine

In the brain slices analysed for antipsychotic-induced *c-fos* activity, we observed consistent tdTomato labelling along the ventricles in Clozapine and Olanzapine treated animals. This labelling was observed in the lateral ventricles as well as the 3^rd^ ventricle (figure 3a–d) and the 4^th^ ventricle (data not shown). Haloperidol and Loxapine showed very minimal labelling along the ventricles (figure 3a–d). Moreover, the cells lining the ventricles were more responsive to Clozapine treatment than the cortical regions that we analysed. These cells showed striking responses even at 5mg/kg of Clozapine (figure 3f).

**Figure 3.**
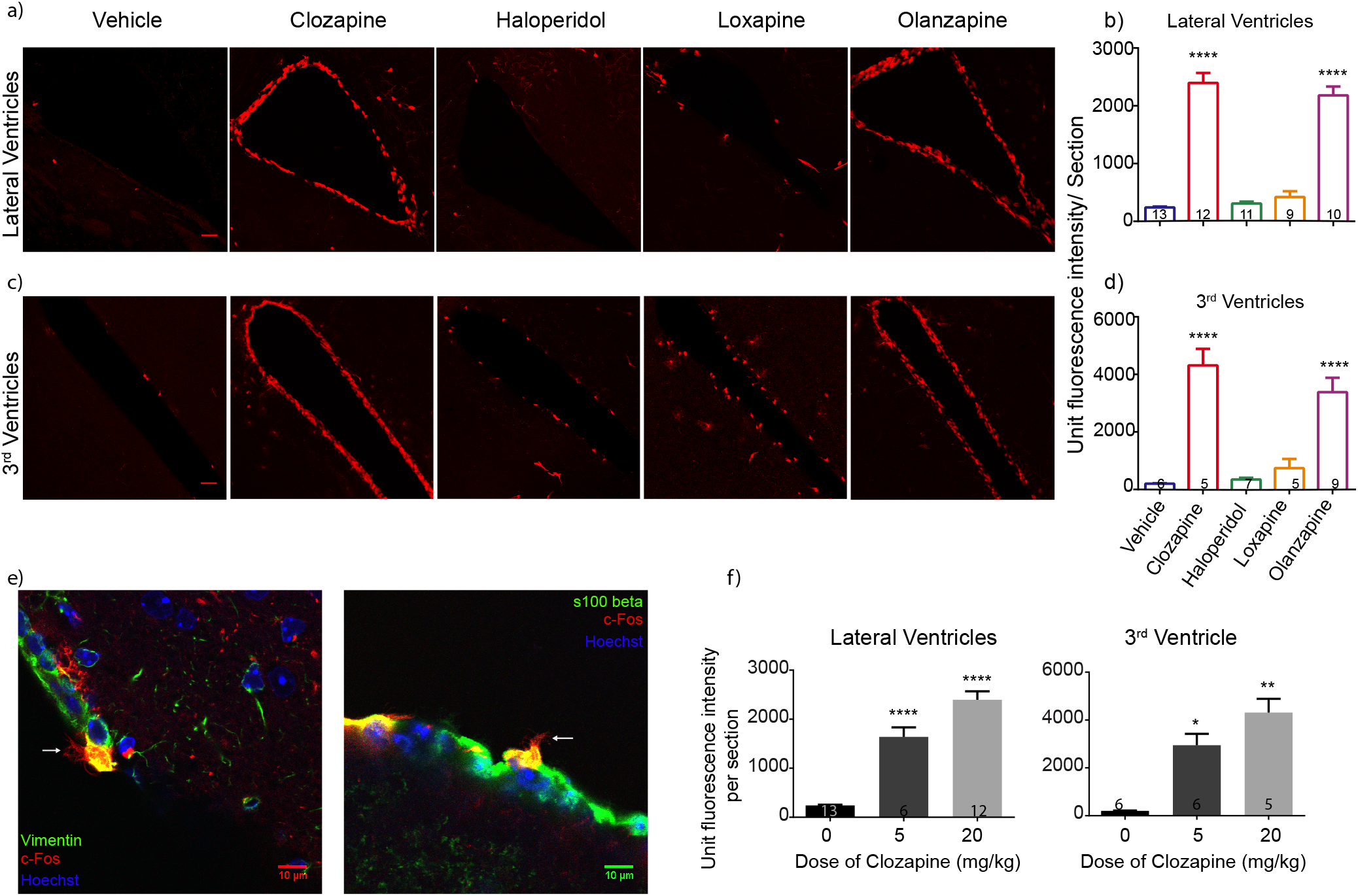
Ependymal cells are the novel cellular target of Clozapine and Olanzapine. a and c) Representative images of the tdTomato positive labelling along the lateral ventricles and the 3^rd^ ventricle, on treatment with various antipsychotics and vehicle. Scale Bar-10 μM. Distinct tdTomato fluorescence can be seen on treatment with Clozapine and Olanzapine. b and d) Quantification of the tdTomato positive labelling along the cells lining the ventricles. Clozapine and Olanzapine showed significantly more fluorescence compared to vehicle. Haloperidol and Loxapine induced *c-fos* activity along the ventricles was comparable to that of the vehicle. Numbers in the bars represent the number of mice in that group. Drug and vehicle groups were compared by one-way ANOVA or Kruskal-Wallis test where appropriate. e) tdTomato positive cells lining the ventricles show multiple cilia and stain positive for the markers of ependymal cells. White arrows point towards ciliated structures. Scale Bar-10 μM f) Graphs represent dose response to Clozapine. Cells lining the ventricles showed significantly higher tdTomato positive labelling even at 5mg/kg dose of Clozapine. one-way ANOVA or Kruskal-Wallis test was performed for data interpretation. Data represented as Mean ± SEM. * p< 0.05, ** p<0.01, *** p< 0.001, **** p< 0.0001.

Atypical antipsychotics have been shown to stimulate neurogenesis in the sub-ventricular zone, the sub-granular zone in hippocampus and in the cortex (Halim et al., 2004; Kodama et al., 2004; Wakade et al., 2002; Wang et al., 2004). Therefore we thought that the tdTomato positive cells lining the ventricles could be the neural stem cells. However, under high magnification these cells showed multiple cilia (Figure 3e), suggesting that these could be the ependymal cells (Brightman and Payal, 1963; Del Bigio, 1995; Jiménez et al., 2014). Additionally, these cells stained positive for the ependymal cell marker Vimentin and S100β (Figure 3e) (Bruni, 1998; Didier et al., 1986; Schnitzer et al., 1981). Therefore we conclude that these were ependymal cells which are novel cellular targets of the antipsychotics-Clozapine and Olanzapine.

### 5-HT_2A_ does not alter the number of tdTomato positive cells in the cortical and thalamic region

5-HT_2A_ has been thought of as an important target for Clozapine’s therapeutic efficacy in mouse models of Schizophrenia (Fribourg et al., 2011; Moreno et al., 2016; Schmid et al., 2014). 5-HT_2A_ expression has also been reported in the various regions of cortex such as piriform cortex, orbital cortex, cingulate cortex, etc. (Miner et al., 2003; Xu and Pandey, 2000). 5-HT_2A_ levels are reported to increase in post-mortem brain (cortical) samples of drug naïve schizophrenic patients (González-Maeso et al., 2008; Muguruza et al., 2013) and chronic treatment with Clozapine reduces levels of 5-HT_2A_ in patients and animal models (Muguruza et al., 2013; Yadav et al., 2011b). Therefore we examined if 5-HT_2A_ receptor would modulate the observed pattern of *c-fos* activity with Clozapine. We crossed the *Htr2a^−/−^* mice, generated by our group (Joshi et al., 2016), into the FosCreER^T2^ (c-Fos Cre ERT2 (B6.129(Cg)-*Fos^tm1.1(cre/ERT2)Luo^*/J) and Lox-tdTomato (B6.Cg-*Gt*(*ROSA*)*26Sor^tm14(CAG-tdTomato)Hze^*/J) background.

We observed that the Clozapine-induced *c-fos* activity in the cortical regions and thalamic regions, was largely unaffected in the *Htr2a^−/−^* background, at both the doses tested-5mg/Kg and 20mg/ (Figure 4a–d). This suggests that 5-HT_2A_ has minimal or no role in determining the number of tdTomato positive neurons in response to Clozapine, in these regions. This result was surprising and suggests that the tdTomato positive cells are the result of complex circuitry and do not rise from direct interactions between 5-HT_2A_ and Clozapine.

**Figure 4.**
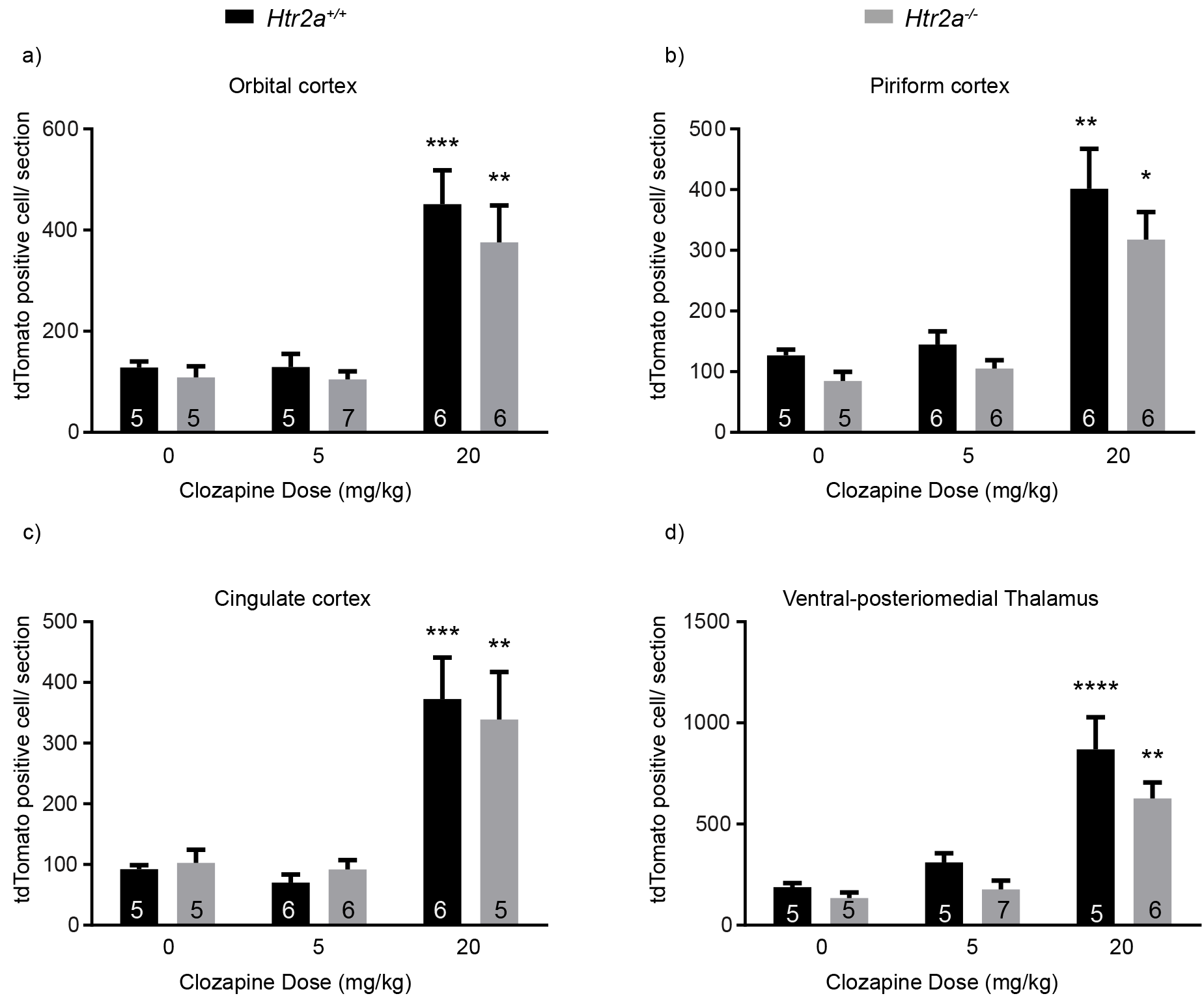
Clozapine induced *c-fos* activity in the cortical regions and Vpm is largely intact in the *Htr2a^−/−^* mice. a- d) *Htr2a^+/+^* and *Htr2a^−/−^* mice do not show differences in the number of *c-fos* positive cells in the orbital cortex, cingulate cortex, piriform cortex and Vpm, at 5 or 20 mg/kg. Numbers in the bars represent the number of mice in that group. two-way ANOVA was performed. Data represented as Mean ± SEM. * comparison between vehicle and treatment for the same genotype. # represents comparison between wild type and mutant under the same conditions. * p< 0.05, ** p<0.01, *** p< 0.001, **** p< 0.0001.

### Clozapine-induced tdTomato labelling in the ependymal cells is modulated by 5-HT_2A_

Unlike the rest of the brain regions, Clozapine-induced tdTomato labelling of ependymal cells was dramatically diminished in the *Htr2a^−/−^* mice even at 5mg/kg of Clozapine (Figure 5a & b). However on increasing the dose to 20mg/kg, labelling of the ependymal cells recovered drastically in the *Htr2a^−/−^* mice, and was statistically indistinguishable from the *Htr2a^+/+^* mice (Figure 5c & d). This data suggests that the genetic deletion of 5-HT_2A_ receptor modulates the Clozapine induced *c-fos* activity in the ependymal cells at low doses of Clozapine.

**Figure 5.**
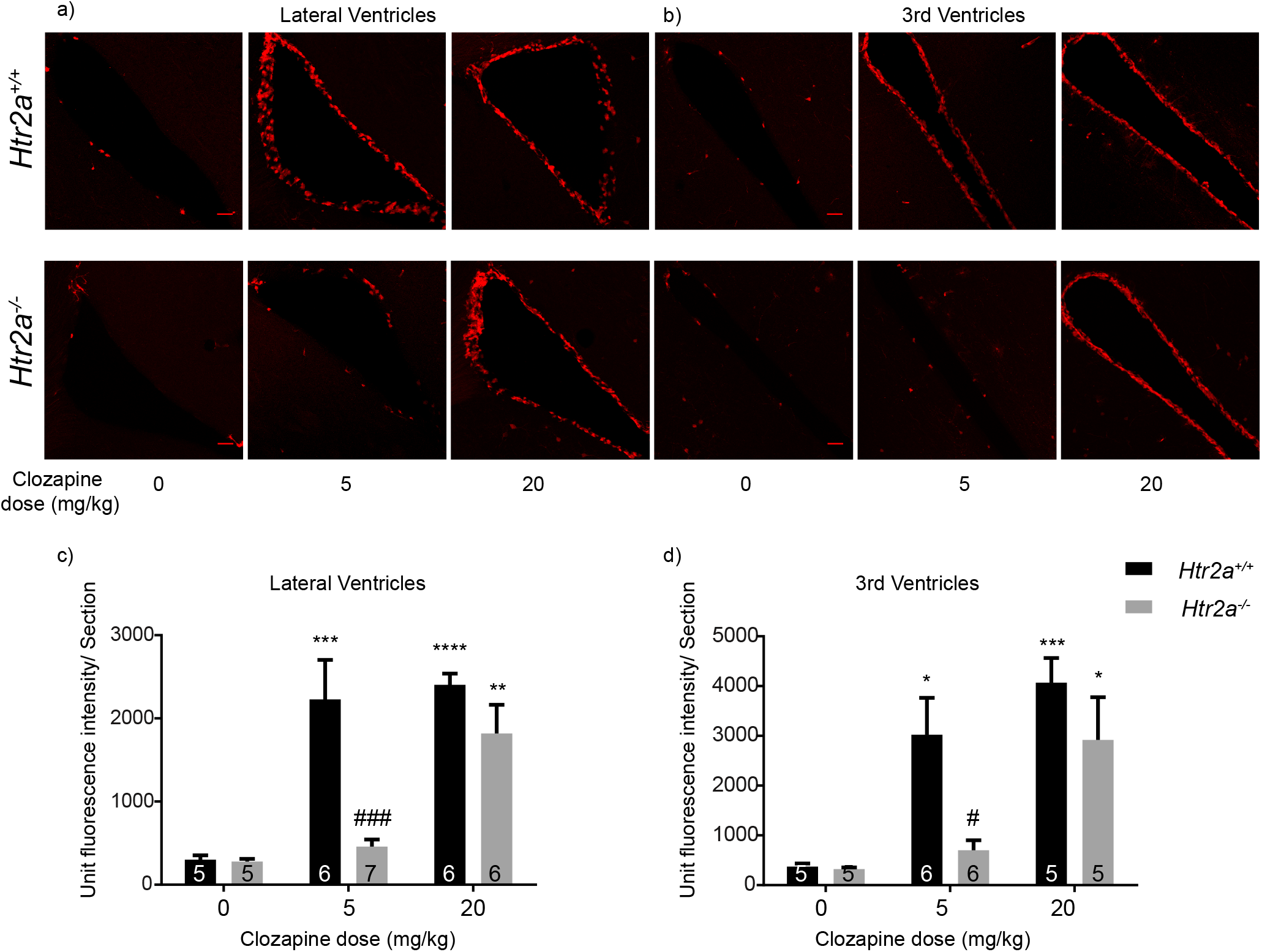
Clozapine induced *c-fos* activity in the ependymal cells is diminished by deletion of 5-HT_2A_. a and b) Representative images of the lateral ventricles and the 3^rd^ ventricles. At 5mg/kg the tdTomato labelling is strikingly diminished in the *Htr2a^−/−^* mice. However at 20mg/kg the activity reappears. Scale Bar-10 μM. c & d) The lateral and the 3^rd^ ventricles showed reduced tdTomato fluorescence at 5mg/kg in the *Htr2a^−/−^* mice compared to the *Htr2a^+/+^*. However at 20mg/kg the fluorescence is statistically not different between the *Htr2a^+/+^* and *Htr2a^−/−^* mice. Numbers in the bars represent the number of mice in that group, two-way ANOVA is performed. Data represented as Mean ± SEM. * comparison between vehicle and treatment for the same genotype. # represents comparison between wild type and mutant under the same conditions. * p< 0.05, ** p<0.01, *** p< 0.001, **** p< 0.0001.

In summary, the FosTrap system has led to the identification of brain areas such as orbital cortex, piriform cortex, Vpm etc. as areas where *c-fos* gets consistently and reproducibly induced by the antipsychotics-Clozapine and Olanzapine. The labelled cells are neuronal and can be potentially accessed ‘live.’ We also observed that ependymal cells up-regulate their *c-fos* activity in response to the antipsychotics Clozapine and Olanzapine. To the best of our knowledge, this is the first report of the effect of antipsychotics on ependymal cells. Furthermore, we observed that absence 5-HT_2A_ did not alter the number of tdTomato positive cells at low or high doses in cortical or thalamic regions that were tested. However it showed a strong dose dependent effect on the tdTomato labelling of ependymal cells.

## Discussion

Thorough understanding of the mechanism of the existing antipsychotics can hold the key to develop safer and sustainable drugs. The pharmacology of the available antipsychotics has been studied quite well, though the neural correlates of these need more investigation. In this study, we have addressed a part of this issue using the FosTrap system devised by Liqun Luo and colleagues.

This system allowed re-examination of the antipsychotic-induced pattern of activity with some distinct advantages. The use of the additional inducible switch using CreER-lox and expression of the fluorescent protein allows for the permanent marking/labeling of the cells, does not require the sacrifice of the animals immediately after the stimulus, allows the examination of living cells and also allows long-term effects of the stimulus to be examined. Therefore FosTrap system can potentially throw different candidates than the conventional techniques. By substituting other proteins and reporters the range and type of experiments can also be extended.

Our experiments have provided novel information as well as corroborated some of the results already available. The Haloperidol induced *c-fos* activity in the striatum has been reported before. This activity is attributed to D2 antagonism and the cataleptic side effect of Haloperidol (Kapur et al., 1995, 2000; Wadenberg et al., 2000). In addition to the typical antipsychotics that we tested (Haloperidol and Loxapine), Olanzapine also showed induction of *c-fos* in the striatum. This was a surprising finding owing to the atypical nature of Olanzapine. In light of this, it would be interesting to assess the catalepsy like behaviour induced by Olanzapine at this dose. However it was difficult to score for catalepsy due to the strong sedation induced by Olanzapine (data not shown).

A parsimonious explanation for the different pattern of antipsychotic-induced activity in the striatum (Figure 1d) can be provided by the relative receptor binding profiles of these drugs. As mentioned above, D2 antagonism is thought to promote catalepsy, whereas 5-HT_2A_ antagonism is thought to improve catalepsy like behaviour (Ansah et al., 2011; Creed-Carson et al., 2011). The balance of affinities for D2 and 5-HT_2A_ may determine the *c-fos* activity in the striatum. Evidently, among the four antipsychotics tested, Haloperidol shows the highest affinity for D2 and Clozapine shows the least. Olanzapine and Loxapine have ten fold more affinity for D2 than Clozapine and more affinity for 5-HT_2A_ than Haloperidol (Roth et a I., 2004).

Clozapine and Olanzapine induced increase in *c-fos* activity in the cortical areas has useful implications, particularly because these regions have been associated with the pathophysiology of Schizophrenia or other mental disorders. For example, reduction in grey matter in the anterior cingulate cortex and orbitofrontal cortex has been observed in postmortem samples of Schizophrenia patients (Fornito et al., 2009; Pantelis et al., 2003). Reduction in the volume of olfactory bulb and deficits in the olfactory capacity have also been reported in patients with Schizophrenia (Moberg et al., 1999; Turetsky et al., 2000). Moreover these cortical areas have been associated with cognitive functions such as reward learning, decision making (Rushworth et al., 2011), working memory (Barbey et al., 2011), etc. *c-fos* activity in the piriform cortex was found to be associated with antidepressant effects (Sibille et al., 1997). Therefore *c-fos* induction in these brain regions may corroborate with the effect of antipsychotics on the negative symptoms or the cognitive symptoms of Schizophrenia.

Antipsychotic-induced increased *c-fos* activity in the Vpm is a novel finding from our work. Hallucinogens such as LSD and DOI have been shown to bring about hallucinogen specific gene regulation in the somatosensory cortex (González-Maeso et al., 2003). Therefore the increased *c-fos* activity in the Vpm, in response to Clozapine and Olanzapine, may highlight the circuitry underlying modulation by antipsychotics in the somatosensory cortex.

Taking into account the high affinity of Clozapine for 5-HT_2A_ and distribution of 5-HT_2A_, we expected Clozapine to show a difference in the number of tdTomato positive cells in the cortical areas. Surprisingly the response of Clozapine was identical in the *Htr2a^−/−^* and *Htr2a^+/+^* background in terms of the number of tdTomato positive cells. It would be of interest to determine the strength of the response and the transcriptome of the ‘tdTomato positive cells’ in the absence of 5-HT_2A_; however, it is beyond the scope of the current article. Also one must remember that the Trap system does not directly report on the intensity of *c-fos* induction in a cell.

In our system we saw *c-fos* activity predominantly in cells with neuronal morphology, except the ependymal cells. Antipsychotic-induced activation of the ependymal cells was an unexpected finding from this study. In Clozapine-induced dose response, most brain areas did not show significant *c-fos* activity at a lower dose of 5mg/kg. However strikingly large number of ependymal cells showed response even at the lower dose of Clozapine i.e., 5mg/kg. This indicates that ependymal cells are more sensitive to Clozapine treatment than the cells in the most of the brain areas that we tested.

Ependymal cells form a barrier between CSF and brain tissue and are involved in diverse functions such as the production and circulation of CSF (Nelson and Wright, 1974; Nielsen et al.; Tissir et al., 2010), transport of water molecules (Jiménez et al., 2014; Nielsen et al.). Also, ependymal cells provide a niche to neural stem cells and promote neurogenesis (Lim et al., 2000; Luo et al., 2008; Tramontin et al., 2003). *c-fos* activity in ependymal cells has been reported in response to formalin-induced acute pain (Palkovits et al., 2007), post hypoxia seizures (Gunn et al., 1990) and on treatment with the antidepressants such as Rolipram (Dragunow and Faull, 1989). Additionally, Serotonin has been shown to increase ciliary beat frequency, which can be blocked by broad spectrum 5-HT2 receptor antagonist-Mianserine (Nguyen et al., 2001). Therefore recruitment of ependymal cells by antipsychotics can provide valuable information into the functioning of antipsychotic drugs and the pathophysiology of mental illness.

Moreover, Clozapine-induced *c-fos* activity in ependymal cells was absent in the *Htr2a^−/−^* mice at 5mg/kg. Interestingly the activity is regained when the dose of Clozapine is increased to 20mg/kg, suggesting that 5-HT_2A_ facilitates the activation of ependymal cells in response to Clozapine and the *Htr2a^−/−^* mice should serve to understand this phenomenon better.

This study has put forth some interesting candidates, but one has to keep certain limitations in mind. Firstly, we have used wild-type or *Htr2a^−/−^* mice, which by themselves are not a model of psychosis. We have so far looked at the acute effects of antipsychotic treatment only. Antipsychotic treatment to patients is often chronic and the therapeutic effects begin to appear later in the treatment whereas side effects appear in the early part of the treatment and may fade out in the later part. Therefore it would be valuable to compare the acute and chronic patterns of antipsychotic activity. Also one must remember that the patterns of activity observed by us can be limited by the differences in the efficiency of ‘TRAP’ in different neural subtypes or brain regions. Variation in efficiency can in part address the lack of antipsychotic induced *c-fos* activity in the nucleus accumbens and medial prefrontal cortex, which have been reported as the targets of antipsychotics earlier.

Taken together, the ‘TRAP’ system has allowed us to identify novel and potentially valuable targets of antipsychotics. This study is a ‘proof of principle’ study showing that the antipsychotic responsive cell populations can be targeted by the TRAP system and made available for further investigation in terms of transcriptome, proteome, metabolome, physiology and even optogenetic modulation of their activity.

## Acknowledgment

We thank Animal Care and Resource centre (ACRC) at the National Centre for Biological Sciences (NCBS) for housing and care of animals. We thank Central Imaging and Flow Cytometry Facility (CIFF) at the NCBS for providing infrastructure for the confocal imaging. We would like to thank Prof. Sanjeev Jain (NIMHANS, Bangalore, India) for discussions and inputs. We thank the funding sources-Department of Biotechnology, Government of India (Grant Number-10961) and intramural funding from the NCBS.

